# Prevalence and trend of malaria with anaemia among under-five children in Jasikan District, Ghana

**DOI:** 10.1101/2020.03.24.005280

**Authors:** William Domechele, George Pokoanti Wak, Francis Bruno Zotor

## Abstract

**Background:** Malaria still remains a major killer of children under-five, claiming the life of one child every two minutes globally. More than 78% of deaths among children under-five in Africa are as a result of malaria infection. Despite the several interventions to reduce malaria and anaemia, the disease remains a global public concern as more children continue to die. This study assessed the prevalence and trend of malaria and anaemia in children under-five years from 2012 to 2016.

**Methods:** We conducted a descriptive cross-sectional study among children under-five with malaria and anaemia who received care at the hospital in Jasikan town, Ghana from 2012 to 2016. Data were extracted from the children’s ward admission and discharge registers. We computed descriptive statistics to describe the data. STATA version 14 was used to carry out the analyses.

**Results:** Out of 30082 malaria cases, 835 were with anaemia from 2012 to 2016. This study demonstrated an overall proportion of malaria with anaemia as 0.028 (28 per 1000 malaria cases). Year 2014 recorded the highest proportion (38 per 1,000 malaria cases) of malaria with anaemia cases in the district. Overall, prevalence rate of malaria and anaemia cases separately was found as 61.5% and 4.4% respectively from 2012-2016. Children within 24-35 months’ age group contributed the highest (28.3%) and 0-11 months accounted for the lowest (12.9%) malaria with anaemia cases. Majority of malaria with anaemia cases 531 (63.6%) occurred in the rainy season from June to July.

**Conclusion:** There is a high prevalence of malaria with anaemia cases among children under-five from 2012-2016. Age and seasonal variation were found to be predictors of an increase in the prevalence of malaria with anaemia.

## Introduction

Malaria and anaemia are frequently co-existing diseases that cause significant morbidity and mortality especially among children-under five years [1]. Studies have shown that children below the age of five years are at the greatest risk of being diagnosed and admitted for malaria infection than any other age groups [2]. The predisposing factors of anaemia in children under-five are diverse, ranging from low immunity, nutritional deficiencies, hookworm infection, HIV, haemoglobinopathies and malaria [3]. In the tropical regions, malaria contributes to high rates (51.4%) of anaemia in young children and it is one of the most crucial factors [3]. A category-specific prevalence study shows 51.4% anaemia as a result of malaria parasites among children under-five [4].

Malaria still remains a major killer of children under-fives, claiming the life of one child every two minutes worldwide [5,6]. Globally, almost half of the world’s population live in malaria endemic areas [7]. Malaria killed an estimated 303,000 children under-five years globally, including 292,000 in the Africa Region between 2010 and 2015 [6].

Malaria accounts for more than 85% in 2008 and 78% in 2013 deaths among children under-five years in Africa [8]. The burden is heaviest in sub-Saharan Africa, where an estimated 90% of all malaria deaths occur in children under-five years, who account for 78% of all deaths [2,3]. In Ghana, malaria is hyper-endemic and accounts for 44% of outpatient attendance, 13% of all hospital deaths, and 22% of mortality among children under-five years of age presenting a serious health problem in Ghana [9]. Anaemia and undernutrition in Rwanda revealed that the prevalence of malaria with anaemia was high among older children within the age-group 24-35 months [10].

A study in Nigeria showed malaria (77.6%) was the commonest condition causing severe anaemia among children under-five (11). Studies in Kenya and Indonesia have revealed anaemia diagnosis status of a child positively associated with malaria inpatients [12,13]. A retrospective study on malaria and associated co-morbidity in children admitted with fever manifestation in Western Ghana revealed the proportion of children with malarial anaemia significantly increased, from 50% in 2010 to 79.2% in 2012 [14]. Increased prevalence of malaria with anaemia (1.6-22.0%) was shown in Papua New Guinea [12,15].

Reduction in malaria and anaemia trends among children under-five has been observed in several studies over a period in Nigeria, Zanziba and Rwanda [16–18]. Even though there was a decline in the prevalence of malaria among children under-five, the number of malaria infection was observed to increase during harmattan and the months of the rainy season, especially September in 2012 and 2013; and June in 2014 [10,16–18]. The number of malaria infection was observed to increase during harmattan and the months of the rainy season, especially in August and September, notwithstanding a decline in the prevalence of malaria among children under-five [16]. Findings from studies in Ejisu-Juaben and Hohoe, Ghana, reported that the prevalence of malaria and anaemia was significantly higher during the pre-rainy season than the post-rainy season [19,20]. The autors elaborated that rainfall increases the possible breading grounds for mosquito larvae, eventually resulting in more vectors to spread the disease during pre-rainy season.

In spite of the above findings elsewhere, little information of this nature is known among children under-five years in the Jasikan District of the Volta Region of Ghana to inform management decision making in the prevention and control of malaria with anaemia among this sub-group. It is therefore, essential to assess the prevalence and trends of malaria with anaemia among this group. This would serve as a source of information for quality assurance and health policy planning in the prevention, control and management of malaria with anaemia among children under-five years.

## Materials and Methods

This assessment was entirely based on primary data from inpatient registers at the hospital. We conducted two in-depth analysis on malaria with anaemia related indicators. Firstly, we determined the proportion of children under-five who had malaria infection and were anaemic between 2012 and 2016. And secondly we assessed the trends in malaria with anaemia cases between 2012 and 2016.

### Study setting

The study was conducted at Jasikan district hospital located in the northern part of the Volta Region of Ghana, which is about 110 kilometres north-east of Ho, the regional capital. The District experiences a double maxima rainfall regime. The major season starts from May and ends in July with its peak in July while the minor season is from September to October with its peak in October [21].

### Study design

The study was a hospital-based cross-sectional study in which patients’ case files and consulting room registers were used to gather information for assessment of the prevalence and trend of malaria with anaemia at the hospital over a period of five years (2012-2016).

#### Study population

The study population were children under-five years of age who attended the hospital between 2012 and 2016.

#### Inclusion and exclusion criteria

All children under-five years of age who attended the hospital and whose records were available for the period of study were included in the study. Children without hospital records or incomplete records were excluded from the study. Also, children under-five years of age without any provisional or definitive diagnosis were excluded.

#### Sample size

All data on malaria and anaemia cases among children under-five in the district hospital for the five-years period was used (January 2012 to December 2016). A total sample size of eight hundred and thirty-five (n=835) children under-five with malaria and anaemia cases were extracted and compiled within the period.

#### Data collection tool

Relevant data were collected from the children’s ward admission and discharge record books using a proforma.

### Data collection

A pre-tested structured compilation sheet was used to retrieve data. Data was collected from relevant registers from January 2012 to December 2016 using a structured compilation sheet to retrieve the data. Criterion approach [22] which is a subset of purposive sampling was used to select all malaria with anaemia cases. Data was collected on the number of children under-five years of age that attended the hospital, month and year of admission, age, sex, diagnosis, and treatment outcome. A child under five years was captured as a case if s/he was diagnosed with malaria as principal and anaemia as additional, resident in any of the communities in the district and within the study period of 2012 to 2016. Data was considered as adequate for each of the years and included in the analysis if information on the indicators selected was available for more than 60% or if less than 30% of consecutive months were missing.

### Data processing and analysis

The data completeness and consistency were checked, variables were coded, and entered into IBM SPSS version 24.0 software. Data analysis was done using STATA version 14.0 (Stata Corp, Texas, USA) on two levels. (1) Analysis of proportions and prevalence: we described the data by computing descriptive statistics, including percentages, frequencies, summary measures, and proportions were used to visualize the data, augmented with graphs drawn using Microsoft Excel program version 2016. (2) Analysis of trends: simple cross-tabulations analysis was used in generating the trends of malaria with anaemia cases within the period of study. Descriptive interpretation was used to understand the nature and pattern of graphs.

### Ethical consideration

Ethical approval for the study was sought and obtained from the Ethical Review Committee of the Ghana Health Service, Ministry of Health (ID NO: GHS-ERC: 71/05/17) with the help of the University of Health and Allied Sciences, School of Public Health, Ho, Volta Region. Also, written permission for the study was obtained from the Jasikan District Health Directorate and the Jasikan District Hospital as well as In-charge of the Health Information Unit. Authorities (the District Health Director and hospital in-charges) were assured of confidentiality.

They were also assured that storage, analysis and reporting of all data including dissemination will be done in formats that will not reveal the identity of patients. After going through the Informed Consent Form, the District Health Director and hospital in-charges were made to indicate their names and signatures in the statement of consent portion to indicate their free will to grant us access to the consulting room registers. Patients privacy was protected by extracting the data in a secluded office.

## Results

### Background characteristics of respondents

We recorded a total of 30082 malaria cases among children under-five from 2012 to 2016. Out of the total cases, 835 had anaemia with the majority being females (50.5%). Age of the cases ranged from 1-58 months, with the mean age of approximately 26 months. Majority (98.4%) of the children under-five years with malaria and anaemia who were admitted were treated and discharged (Table 1).

**Table 1:** Demographic characteristics and treatment outcome among children under-five.

| Variable | Frequency [N=835] | Percentage (%) |
| --- | --- | --- |
| **Age group (months)** |  |  |
| 0-11 | 108 | 12.9 |
| 12-23 | 191 | 22.9 |
| 24-35 | 236 | 28.3 |
| 36-45 | 189 | 22.6 |
| 46-59 | 111 | 13.3 |
| **Sex** |  |  |
| Male | 413 | 49.5 |
| Female | 422 | 50.5 |
| **Treatment outcome** |  |  |
| Treated | 822 | 98.4 |
| Died | 13 | 1.6 |

### Proportion of children under-five having malaria infection and are anaemic

As shown in Table 2, the overall proportion of malaria with anaemia cases among children under-five from 2012-2016 was 28 per 1,000 malaria cases. The highest proportion (38 per 1,000) of malaria with anaemia cases was recorded in 2014. The prevalence of malaria and anaemia cases separately among the children ranged from approximately 58.7% to 62.7% and 2.6% to 6.7% respectively. The highest (62.7%) prevalence of malaria cases was observed in both Year 2015 and 2016 while that of anaemia was 6.7% in the year 2016. The overall prevalence for malaria among all the children was 61.5% whereas that of anaemia was 4.4%.

**Table 2:** Yearly prevalence rate and proportion of malaria with anaemia among children under-five in Jasikan District from 2012-2016.

| Year | Population at risk | Malaria Cases |  | Anaemia Cases |  | Malaria with Anaemia Cases |  |
| --- | --- | --- | --- | --- | --- | --- | --- |
|  |  | Total (N) | Prevalence (%) | Total (N) | Prevalence (%) | Total (N) | Proportion Per 1000 |
| 2012 | 9,517 | 5,861 | 61.6 | 505 | 5.3 | 144 | 25 |
| 2013 | 9,649 | 5,663 | 58.7 | 260 | 2.7 | 189 | 33 |
| 2014 | 9,781 | 6,037 | 61.7 | 464 | 4.7 | 232 | 38 |
| 2015 | 9,913 | 6,220 | 62.7 | 255 | 2.6 | 126 | 20 |
| 2016 | 10,043 | 6,301 | 62.7 | 670 | 6.7 | 144 | 23 |
| **Total** | **48,903** | **30,082** | **61.5** | **2,154** | **4.4** | **835** | **28** |

### Trend of children under-five with malaria and anaemia cases from 2012 to 2016

#### Trends in malaria and anaemia

Fig 1 depicts a secular trend of malaria with anaemia among children under-five from 2012-2016. The pattern of malaria with anaemia cases appeared to be on a rise and fall recapping schema. It fluctuated with an unclear direction of how it moved. Malaria with anaemia increased continuously from 2012 at 144 cases and peaked in 2014 to 232 cases. Thereafter, there was a sharp decline from 232 to 126 cases in 2015, then increased again fom 126 to 144 cases in 2016. Both sexes in 2014 recorded almost the same number of cases. Between 2015 and 2016 cases for both males and females increased slightly.

**Fig 1:**
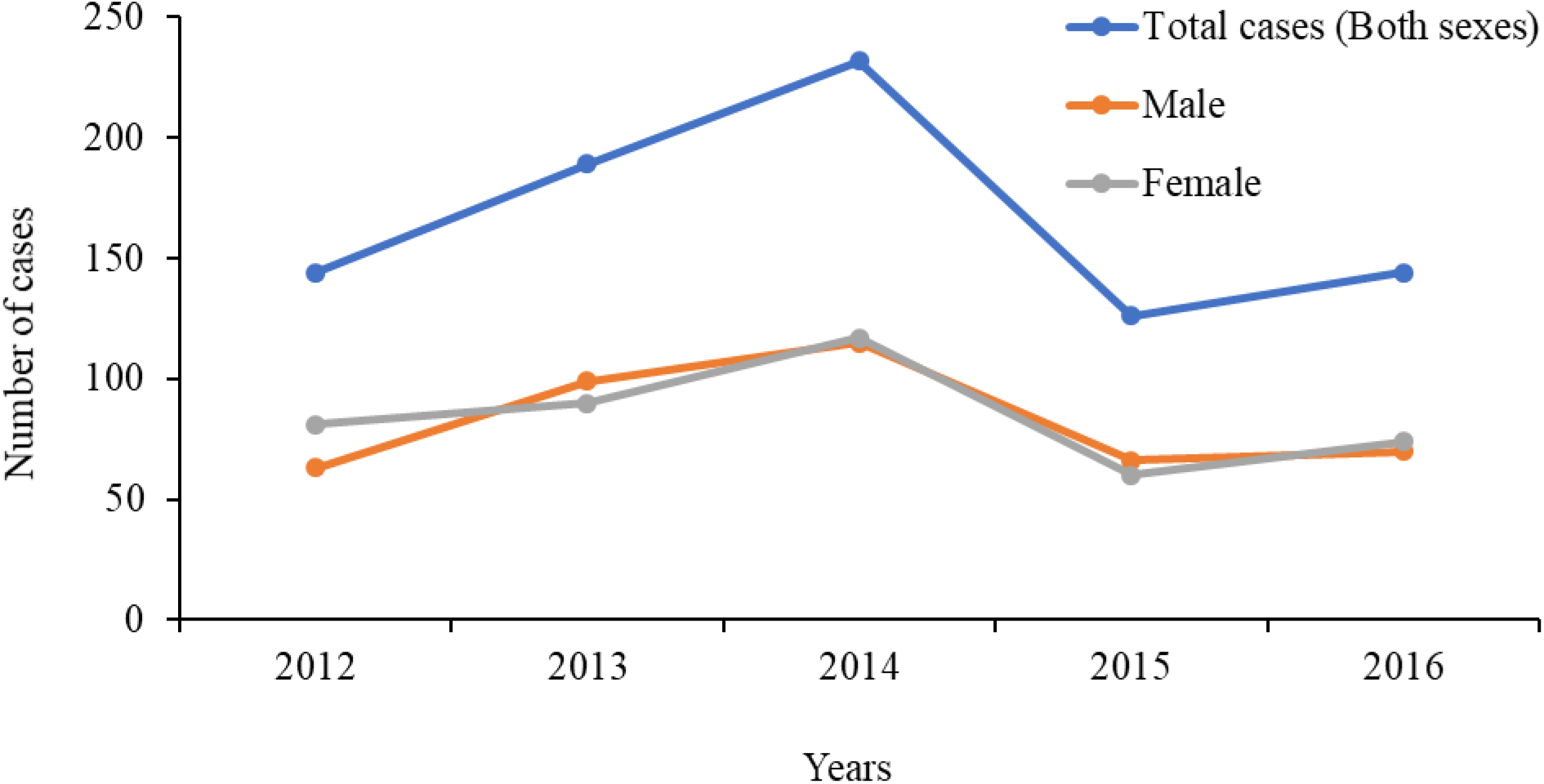
Trend of malaria with anaemia among children under five in Jasikan District from 2012-2016.

#### Seasonal trend in malaria with anaemia cases

Fig 2 depicts seasonal trend of malaria with anaemia cases. There was a rise and fall pattern of malaria with anaemia cases in the harmattan season (November to April). Majority of the cases that were recorded in 2012, 2013 and 2015 in the harmattan season peaked in December and decreased in January; then increased sharply from April to May except year 2015. The pattern of rainy season (June to October) averagely had higher peaks of cases from May to September within all the years. Rainy season for 2013 and 2014 peak occurred in July which corresponds to the general rainy season.

**Fig 2:**
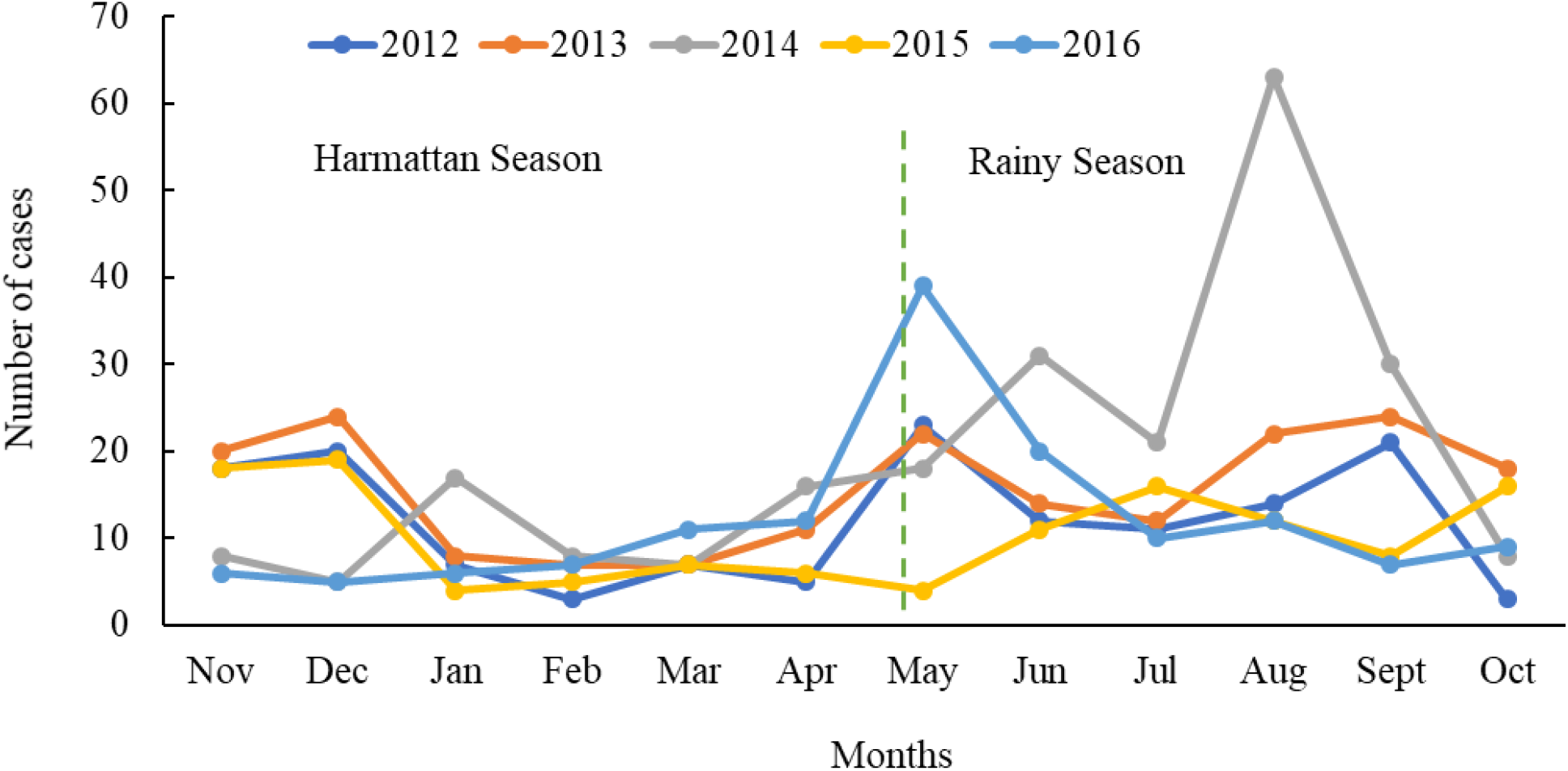
Seasonal trend of malaria with anaemia among under-five children in Jasikan District from 2012-2016.

#### Seasonal pattern of malaria with anaemia cases

There was no clear seasonal pattern of malaria with anaemia cases within the period 2012-2016. However, there was a rise and fall repetition of malaria with anaemia cases as shown in Fig 3. Harmattan season had a steady increase of prevalence of malaria with anaemia among children under-five in the month of December for all the years which clearly depict a peak within the period under study except 2014 and 2016. The number of cases declined slowly after the peak in 2013 until in another harmattan season where it dropped sharply in the month of December in the year 2014. There was always a decline in the number of cases in every one or two months to the end of every harmattan (May) season except in 2015 where the cases decreased continuously from March to May. The pattern of the rainy season appeared to be increasing from August except 2015. A peak for rainy season was observed in the month of August which was higher than that of the harmattan season.

**Fig 3:**
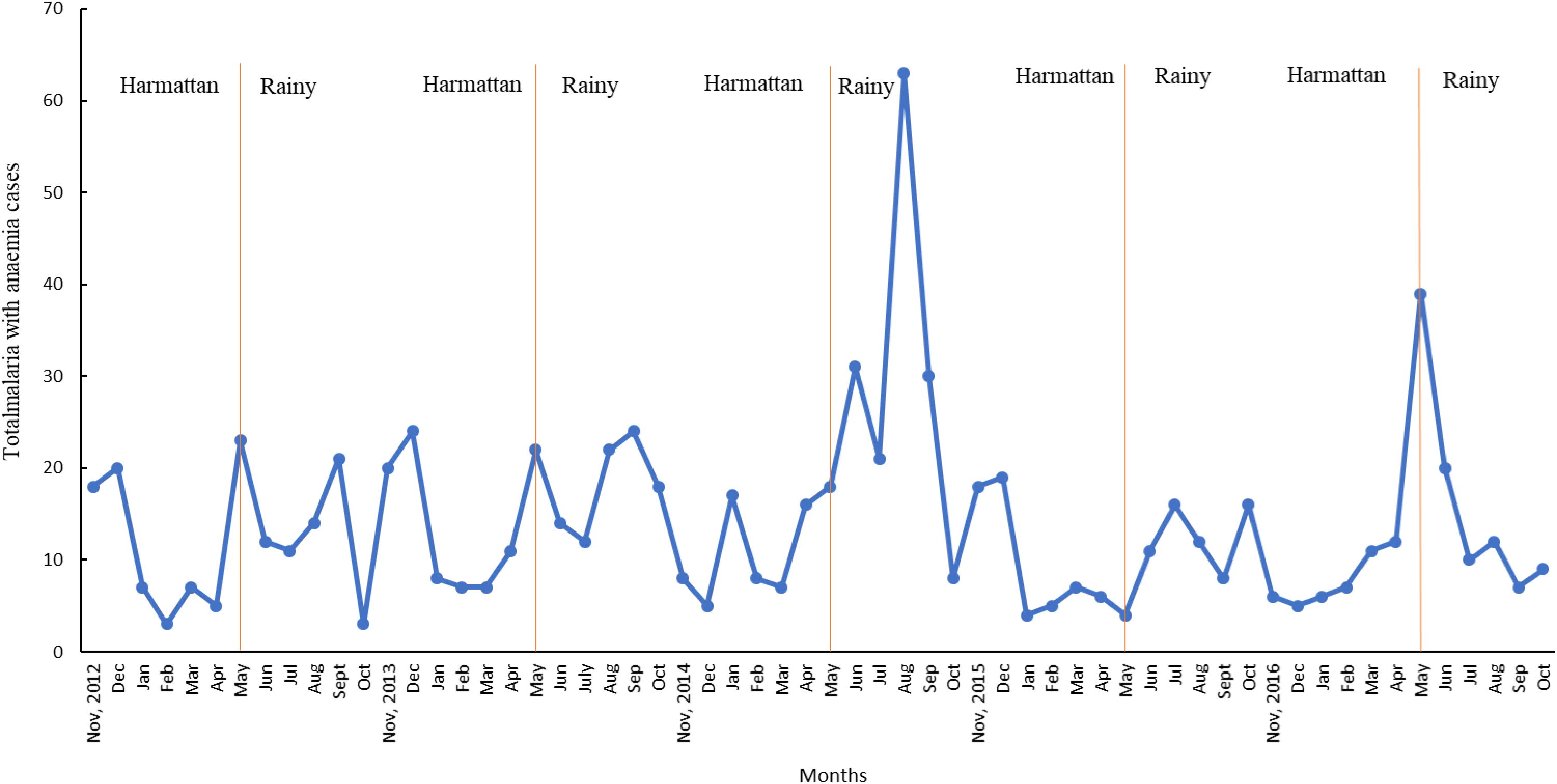
Seasonal pattern of malaria with anaemia cases among under-five children in Jasikan District from 2012-2016.

## Discussion

The purpose of this study was to assess the prevalence and trends of malaria with anaemia among children under-five in the Jasikan District of the Volta region of Ghana. Regarding age distribution, a larger percentage (28.3%) with malaria and anaemia were within the age group of 24 - 35, followed by 12 - 23 months (22.9%) and 36 - 45 (22.6%). This may imply that the increased prevalence of malaria with anaemia in children aged 24-35 months in this study may be an indication of persistent anaemia after treatment of malaria due to prolonged weak immune response. These findings corroborate with a study done in Rwanda [1]. On the contrary, a retrospective study conducted by Orish [14] in Sekondi-Takoradi of Western Ghana on anaemia and associated co-morbidity in children found a significant increase in admission of children younger than five years of age, with a marked increase seen in those younger than one year of age over the three-year study period.

Furthermore, the study showed there was a slight female preponderance of malaria infection with anaemia. The gap could depend on a complex mix of biological factors (immune system, genes and chromosomes, hormones, reproductive anatomy, metabolism). On the contrary, findings from a study reported when it comes to health, males are the weaker sex throughout life [23]. This study findings differ from a study carried out in Anambra State, Nigeria by Chineke et al and Kunihya et al where males have a higher prevalence of malaria with anaemia than females [16,24]. In the same line, another study findings in Benin City, Nigeria found higher malaria prevalence in males (57%) than in females (43%) although there was no significant association between gender and malaria [25]. But why slightly higher prevalence of malaria with anaemia among children under-five females found in this study? It is the $64,000 question; hence further research needs to be carried on to investigate the contradiction.

Malaria burden in Jasikan district, as in most parts of Ghana, has remained high and in many areas even heightened over the years. Significant reductions in anaemia prevalence have been achieved or recorded following malaria control programs in endemic areas. Low prevalence of anaemia found in this study differs with that of Kiggundu et al. (2013) who reported a high prevalence of anaemia (56.3%) among children under-five in Southwestern Uganda [26]. Furthermore, this finding contrasts a higher prevalence of anaemia among children under-five years in Navrongo, Northern part of Ghana [10]. This discrepancy could be as a result of variations in the number of years studied including geographical location.

This study found that in general, 28 out of every 1,000 children under-five having malaria infection are anaemic, and variations were found between five years (20-38 per 1000 malaria cases). This goes to support the results of a prospective study conducted in Papua New Guinea. It was shown that the prevalence of malaria with anaemia among children under-five years of age was between 1.6-22.0% [27,28]. Notwithstanding, this finding disagrees with a study conducted in Western Ghana as they reported a low proportion (20.8%) of malaria with anaemia cases among children under-five [29]. The Jasikan District situated in the Volta Region of Ghana is located in the malaria-endemic zone and might have contributed to the increased proportion of malaria with anaemia cases as compared to Western Ghana. On the contrary, this finding is comparatively lower than the 76.8% prevalence reported in children under-five years from a survey conducted in Southwestern Uganda [26]. Higher prevalence reported in Uganda could be attributed to multifactorial causes such as hookworm and nutritional deficiencies including the location of the district in a malaria-endemic zone.

Trends in the number of malaria with anaemia cases among children under-five can be a powerful means to measure the effectiveness of malaria and anaemia interventions. Hence, regular trend analysis regarding malaria with anaemia among children under-five years is very crucial for planning and prevention. This study has shown that malaria with anaemia cases is more common during the rainy season, as in the month of August which recorded the highest prevalence of infection in 2013 and 2014.

In addition, there was a peak prevalence of malaria with anaemia cases in every August to September of the five years studied. On the contrary, almost all the least number of cases were recorded in every February This result is comparable with the findings of similar studies in Ejisu-Juaben of the Ashanti Region of Ghana which revealed that malaria with anaemia prevalence was highest during the rainy season [10,19,30,31]. The occurenece of malaria transmission among children under-five throughout the year has showed marked seasonal influences on infected children to develop anaemia.

High peak of malaria with anaemia cases in August and September could be due to heavy rainfall during that time of the year as well as environmental factors such as improper environmental management. Presence of stagnant water during the rainy season provides breeding sites for mosquitoes which might have resulted in the high prevalence of malaria with anaemia cases [32]. Inability to control for behavioural changes towards malaria and anaemia awareness among caretakers could also be a contributory factor.

The study found that the observed trend of malaria with anaemia cases was found to have peaked continuously from 2012 to 2014. This finding is in contrast with a study done in Nigeria. Chineke et al (2016) found the prevalence of malaria and anaemia trend among children under-five in 2012, 2013 and 2014 to be 77.0%, 72.4% and 60.8% respectively which depicts a progressive decline in the proportion of malaria with anaemia cases [16].

## Conclusion

Results from this study have shown that the proportion of malaria with anaemia was high among children under-five in the Jasikan District of the Volta region of Ghana. This was associated with the rainy season and demographic characteristics such as age and sex. Rainy season is the period with the highest malaria infection with anaemia.

### Limitation of the study

Missing and incomplete information in the registers were the most important limitations of the study. However, each source of data was used to compliment the other under such a difficult circumstance.

## Acknowledegments

We acknowledge the District Director of Health Services and people of the study communities as well as management of the Jasikan District Hospital and Health Directorate for providing us with the needed support during the data collection period upon which the findings of this study were based.

